# Micro-Mel and Mini-Mel short life cycle dwarf lines with *Solanum anguivi* introgressions as the first model varieties for eggplant research and breeding

**DOI:** 10.1101/2025.03.07.642017

**Authors:** Marina Martínez-López, Virginia Baraja-Fonseca, Edgar García-Fortea, Mariola Plazas, Santiago Vilanova, Jaime Prohens, Pietro Gramazio

**Affiliations:** Instituto de Conservación y Mejora de la Agrodiversidad Valenciana, Universitat Politècnica de València, Camino de Vera 14, 46022 Valencia, Spain; Seeds for Innovation, Parque Científico-Tecnológico de Almería, Calle Regaliz, 6, 04007, Almería, Spain

**Keywords:** Micro-Mel, Mini-Mel, dwarfism, model plant, *Solanum melongena*, phenotyping, regeneration

## Abstract

Eggplant (*Solanum melongena*) research has been hindered by the lack of suitable model plants. The development of Micro-Mel and Mini-Mel, two short-cycle dwarf model lines with introgressions from the wild relative *S. anguivi*, represents a breakthrough in eggplant research. Both lines were characterized with 30 phenotypic descriptors and compared to their parental lines and the Micro-Tom tomato. Micro-Mel has determinate growth, early flowering (41.0 days post-transplantation), and physiological fruit maturity at 103.7 days. Mini-Mel exhibits indeterminate growth, later flowering (57.2 days) and fruit maturity (114.7 days), with a more conventional architecture. Micro-Mel and Mini-Mel complete 3.5 and 3.0 full seed-to-seed cycles per year, respectively. Their compact size makes them ideal for space-limited environments, controlled growth chambers, speed breeding, ornamental horticulture, and urban horticulture. To validate their potential for gene editing, *in vitro* regeneration experiments were performed on a medium containing zeatin riboside. Regeneration was similar to parental *S. melongena* (1.2 shoots/explant), with Micro-Mel developing 1.0 and Mini-Mel 0.8. Whole-genome sequencing (23X coverage) identified 179,653 SNPs between the parental lines, revealing heterozygosity rates of 0.00098% for Micro-Mel and 0.00123% in Mini-Mel. Approximately 8.50% and 10.79% of their genomes were introgressed from *S. anguivi*, respectively. Analysis identified 18 shared introgressions from *S. anguivi* among Micro-Mel, Mini-Mel, and four backcross dwarf materials derived from the same original plant. Notably, three orthologues of genes associated with dwarfism in tomato (*SlDREB1, SlER*, and *SlSERK1*) were identified within these introgressions. Micro-Mel and Mini-Mel represent transformative tools for eggplant research, accelerating genetic, physiological, and breeding studies.

## 1. Introduction

Eggplant (*Solanum melongena* L., 2n = 2x = 24), also known as aubergine or brinjal, is native to Southeast Asia and is congeneric to other major vegetable crops such as tomato (*S. lycopersicum* L.) and potato (*S. tuberosum* L.). Globally, eggplant ranks fourth in vegetable production after tomato, watermelon, and cucumber, with a total output of 60.8 million tons in 2023 [1]. Despite its economic importance, research and breeding in eggplant lag behind other major Solanaceae horticultural crops, such as tomato and pepper, hindering substantial genetic and agronomic advancements [2]. Although eggplant displays considerable morphoagronomic diversity [3], no dwarf fast-cycle materials have been identified or developed as model systems for research. Such materials could enable rapid genetic studies, accelerate breeding programs, and enhance phenotypic and physiological evaluations, as demonstrated in other major crops [4].

One of the most notable examples of interest in dwarf materials is found in tomato, where Scott and Harbaugh (1989) developed the dwarf variety ‘Micro-Tom’ by crossing the cultivars Florida Basket and Ohio 4013-3 Scott and Harbaugh (1989). The dwarf phenotype of Micro-Tom is derived from mutations in, at least, the *DWARF* (D) and *SELF PRUNING* (SP) genes [6]. The compact size of Micro-Tom (10-20 cm), its rapid cycle (70-90 days from sowing to fruit maturity), and its high efficiency in genetic transformation make it an exceptional model for tomato research, allowing four to five generations per year in small spaces with high plant density [4,7–9].

Micro-Tom has been extensively employed in tomato research. Among its many applications, Micro-Tom has facilitated the development of Near-Isogenic Lines (NILs) to enhance genomic studies and the production of parthenocarpic plants using CRISPR/Cas9 technology, demonstrating its utility in genetic engineering [10]. Also, Micro-Tom has been instrumental in the identification of Quantitative Trait Loci (QTLs) associated with fruit weight [11], as well as in studies addressing cold tolerance [12] and salt stress [13]. Additionally, the complete genome assembly of Micro-Tom has been released, providing a valuable resource for genomic studies [14]. Recently, the genomic variation within different Micro-Tom lines has been studied by performing whole-genome sequencing [15].

Beyond tomato, among vegetable Solanaceae crops, the dwarf Micro-Pep variety of pepper (*Capsicum annuum* L.), characterized by short internodes, whorled phyllotaxy, lanceolate leaves, and attenuated rounded pungent fruits, has been developed [16,17]. Two versions of Micro-Pep contrasting for fruit colour, Micro-Pep Yellow and Micro-Pep Red, facilitated the genetic dissection of fruit colour [16]. Likewise Micro-Tom, Micro-Pep is also used as an ornamental variety [17].

In the *Brassicaceae* family, a spontaneous dwarf mutant of cabbage (*Brassica oleracea* L.) was identified, displaying a distinct phenotype characterized by reduced plant height, wrinkled leaves, a non-heading growth habit, and significantly lower self-fertility compared to the wild type [18]. Comprehensive analyses encompassing phenotypic characterization, transcriptomics, and phytohormone profiling were undertaken to elucidate the genetic and molecular basis of the observed dwarfism. The findings revealed that the dwarf phenotype resulted from mutations in genes involved in cell division and phytohormone signalling pathways [18].

In addition to a small size and short life cycle, a model plant should ideally display robust *in vitro* regeneration to facilitate biotechnological tools such as *Agrobacterium*-mediated transformation and gene editing. In this context, Micro-Tom has been widely used for the functional validation of candidate genes, including the *Rg1* allele for improved *in vitro* regeneration [19], *SlDELLA* for the regulation of gibberellic acid signalling [20], and *SlWRKY29* for enhanced disease resistance and pattern-triggered immunity [21]. However, unlike tomato, eggplant exhibits partial recalcitrance to *in vitro* culture, posing significant challenges for biotechnological applications [22,23]. The limited number of available organogenesis protocols typically relies on various combinations and concentrations of phytohormones to induce regeneration, often with inconsistent success [23–25]. Despite these challenges, a handful of gene editing and transformation protocols have been developed, indicating that progress is being made in overcoming the recalcitrance of eggplant [26–28].

In addition to their research applications, dwarf plants can also offer significant ornamental value. Dwarf cultivars in various species, such as dwarf tomatoes [29–31], compact peppers [32,33], and miniature cabbages [34–36] are widely acknowledged for their ornamental appeal and suitability for small-scale gardens and indoor cultivation.

This study presents the development of two compact eggplant model varieties, Micro-Mel and Mini-Mel, derived from interspecific hybridization between *S. melongena* and its wild relative, *S. anguivi* Lam. Micro-Mel and Mini-Mel exhibit phenotypic and phenological characteristics that establish them as model plants for eggplant research, comparable to the widely adopted Micro-Tom model in tomato. Micro-Mel has a dwarf plant size, short internodes, rapid development, multiple inflorescences, and numerous small white fruits. In contrast, Mini-Mel displays a compact size but with longer internodes, greater height, and slightly slower development compared to Micro-Mel, bridging the latter with a more typical standard eggplant. This more conventional plant architecture makes Mini-Mel particularly advantageous for physiological studies, modelling normal growth patterns, but requiring less time and space and allowing for higher plant density than standard varieties. The combination of these traits renders both Micro-Mel and Mini-Mel highly suitable for cultivation in small pots, facilitating their growth under controlled conditions in climatic chambers, as well as in greenhouses.

In this study, we perform a comprehensive phenotypic comparison of Micro-Mel and Mini-Mel with their parental lines and Micro-Tom, focusing on the phenotypic characteristics that distinguish these varieties. Regeneration experiments were conducted to assess the feasibility and potential of Micro-Mel and Mini-Mel as model plants for *in vitro* culture experiments. Additionally, we performed genomic characterization of the two compact eggplant model varieties to identify potential introgressions and genomic regions associated with their distinctive phenotypes. By developing Micro-Mel and Mini-Mel as model eggplant varieties, this study seeks to establish a robust platform for genetic and phenotypic studies in this crop, addressing a significant gap in eggplant research and breeding programs.

## 2. Results

### 2.1 Phenotyping of Micro-Mel and Mini-Mel

The development of Mini-Mel and Micro-Mel resulted in highly stable and compact lines under diverse environmental conditions. Variety fact sheets providing detailed information on Micro-Mel and Mini-Mel, along with images of the plants, flowers, and fruits, are included in Supplementary Material (Figure S1 and S2).

#### 2.1.1 Vegetative traits

At all times points, except 30 days post-transplantation (30 d), Micro-Mel and Mini-Mel exhibited significantly shorter plant heights (P-height) compared to the other three evaluated genotypes (ANG1, MEL1, and Micro-Tom) (Figure 1, Table 1). At 30 d, Micro-Mel was slightly taller than Mini-Mel. However, this trend reversed over time, with Mini-Mel significantly surpassing Micro-Mel in height by 90 d post-transplantation. Micro-Mel reached its maximum height at 60 d, whereas Mini-Mel continued growing until 90 d before attaining its final height. Micro-Tom consistently displayed the greatest height at earlier stages, although it was not significantly different from ANG1 and MEL1 at later stages (Figure 1, Table 1).

**Table 1.**
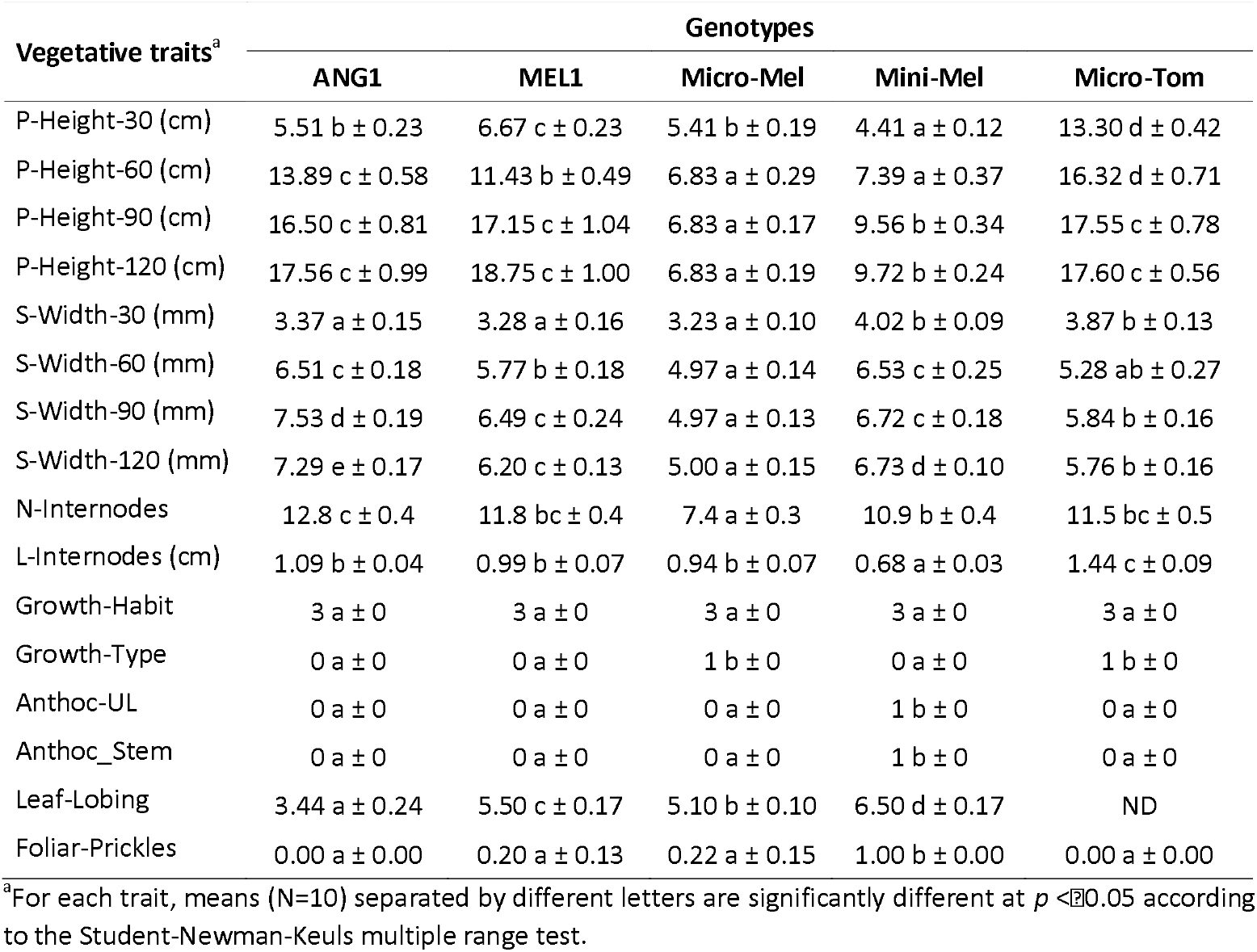
Mean values (± standard error) of the vegetative traits phenotyped for *S. anguivi* ANG1 and *S. melongena* MEL1 parental lines, Micro-Mel, Mini-Mel eggplant model varieties, and Micro-Tom tomato model variety. Ten plants were evaluated for each genotype.

| Vegetative traits<sup>a</sup> | Genotypes |  |  |  |  |
| --- | --- | --- | --- | --- | --- |
|  | ANG1 | MEL1 | Micro-Mel | Mini-Mel | Micro-Tom |
| P-Height-30 (cm) | 5.51 b ± 0.23 | 6.67 c ± 0.23 | 5.41 b ± 0.19 | 4.41 a ± 0.12 | 13.30 d ± 0.42 |
| P-Height-60 (cm) | 13.89 c ± 0.58 | 11.43 b ± 0.49 | 6.83 a ± 0.29 | 7.39 a ± 0.37 | 16.32 d ± 0.71 |
| P-Height-90 (cm) | 16.50 c ± 0.81 | 17.15 c ± 1.04 | 6.83 a ± 0.17 | 9.56 b ± 0.34 | 17.55 c ± 0.78 |
| P-Height-120 (cm) | 17.56 c ± 0.99 | 18.75 c ± 1.00 | 6.83 a ± 0.19 | 9.72 b ± 0.24 | 17.60 c ± 0.56 |
| S-Width-30 (mm) | 3.37 a ± 0.15 | 3.28 a ± 0.16 | 3.23 a ± 0.10 | 4.02 b ± 0.09 | 3.87 b ± 0.13 |
| S-Width-60 (mm) | 6.51 c ± 0.18 | 5.77 b ± 0.18 | 4.97 a ± 0.14 | 6.53 c ± 0.25 | 5.28 ab ± 0.27 |
| S-Width-90 (mm) | 7.53 d ± 0.19 | 6.49 c ± 0.24 | 4.97 a ± 0.13 | 6.72 c ± 0.18 | 5.84 b ± 0.16 |
| S-Width-120 (mm) | 7.29 e ± 0.17 | 6.20 c ± 0.13 | 5.00 a ± 0.15 | 6.73 d ± 0.10 | 5.76 b ± 0.16 |
| N-Internodes | 12.8 c ± 0.4 | 11.8 bc ± 0.4 | 7.4 a ± 0.3 | 10.9 b ± 0.4 | 11.5 bc ± 0.5 |
| L-Internodes (cm) | 1.09 b ± 0.04 | 0.99 b ± 0.07 | 0.94 b ± 0.07 | 0.68 a ± 0.03 | 1.44 c ± 0.09 |
| Growth-Habit | 3 a ± 0 | 3 a ± 0 | 3 a ± 0 | 3 a ± 0 | 3 a ± 0 |
| Growth-Type | 0 a ± 0 | 0 a ± 0 | 1 b ± 0 | 0 a ± 0 | 1 b ± 0 |
| Anthoc-UL | 0 a ± 0 | 0 a ± 0 | 0 a ± 0 | 1 b ± 0 | 0 a ± 0 |
| Anthoc_Stem | 0 a ± 0 | 0 a ± 0 | 0 a ± 0 | 1 b ± 0 | 0 a ± 0 |
| Leaf-Lobing | 3.44 a ± 0.24 | 5.50 c ± 0.17 | 5.10 b ± 0.10 | 6.50 d ± 0.17 | ND |
| Foliar-Prickles | 0.00 a ± 0.00 | 0.20 a ± 0.13 | 0.22 a ± 0.15 | 1.00 b ± 0.00 | 0.00 a ± 0.00 |

<sup>a</sup>For each trait, means (N=10) separated by different letters are significantly different at $p < 0.05$ according to the Student-Newman-Keuls multiple range test.

**Figure 1.**
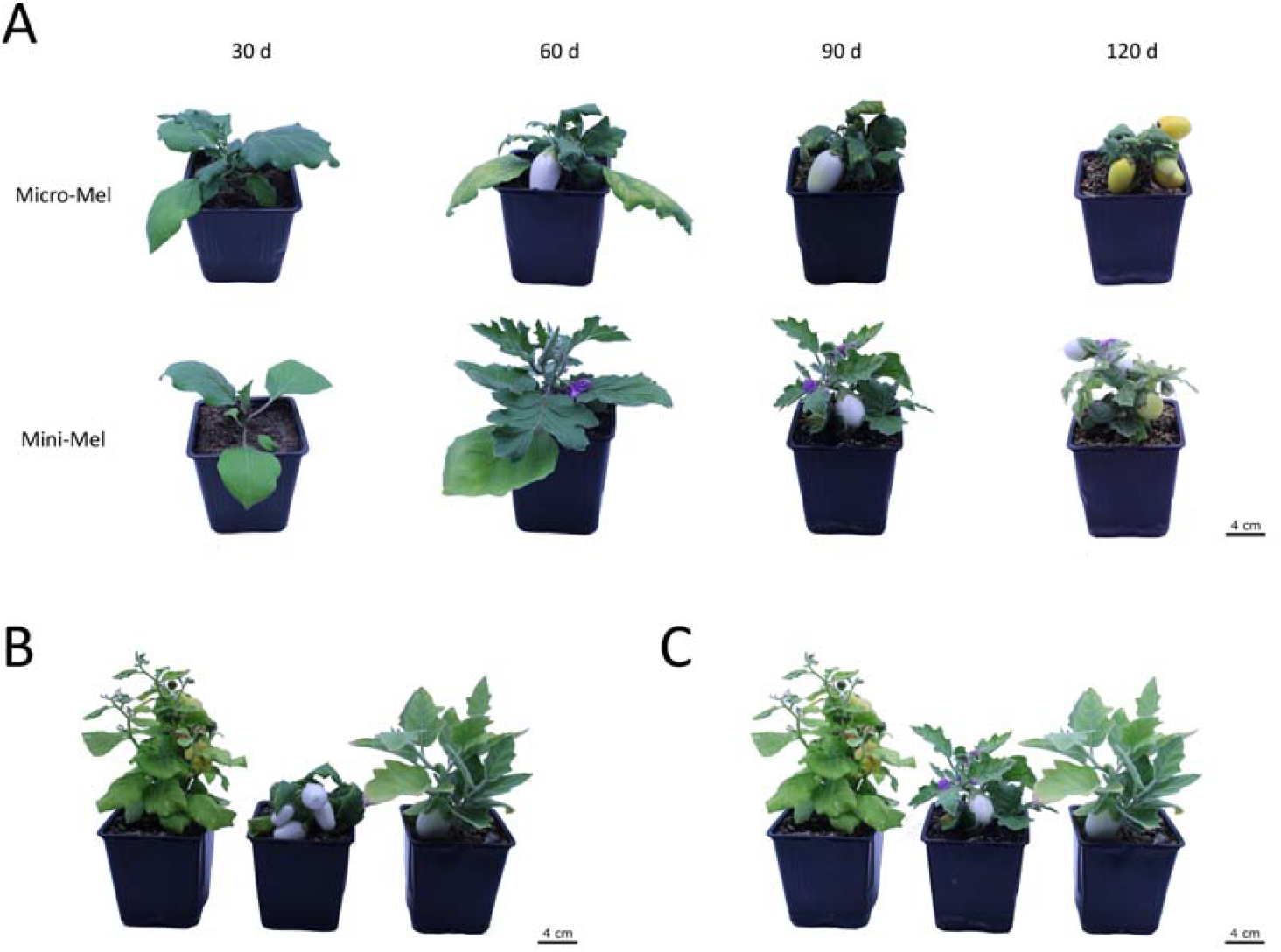
**(A)** Developmental progression of Micro-Mel and Mini-Mel from 30 to 120 days post-transplantation. Top: Micro-Mel. Down: Mini-Mel. From left to right: 30, 60, 90, and 120 days post-transplantation. Comparison of Micro-Mel (**B**, center) and Mini-Mel (**C**, center) with their parental lines ANG1 (left) and MEL1 (right) at 90 days post-transplantation.

At 30 d post-transplantation, Mini-Mel and Micro-Tom exhibited the thickest stems among the genotypes (S-Width-30). By 60 d (S-Width-60), Micro-Mel had the narrowest stem width, comparable to Micro-Tom, while ANG1 and Mini-Mel displayed the thickest stems. From 90 d onward (S-Width-90/120), Micro-Mel consistently had the thinnest stems, whereas ANG1 had the thickest. No significant variations in stem width were observed among the genotypes beyond 90 d (Table 1).

Regarding the number of internodes (N-Internodes), Micro-Mel displayed the fewest, followed by Mini-Mel, which was not significantly different from MEL1, Micro-Tom and ANG1. In terms of internode length (L-Internodes), Mini-Mel had the shortest internodes, while Micro-Tom exhibited the longest. No significant differences were observed among ANG1, MEL1, and Micro-Mel for internode length (Table 1). The growth habit (Growth-Habit) was upright in all the genotypes, but for growth type (Growth-Type), both Micro-Mel and Micro-Tom had determinate growth, while Mini-Mel had indeterminate growth similar to the parental lines ANG1 and MEL1. Anthocyanin presence was observed only in Mini-Mel, occurring in both upper leaves (Anthoc-UL) and stems (Anthoc-Stem). Mini-Mel also had significantly stronger leaf blade lobing (Leaf-Lobing) and higher foliar prickliness (Foliar-Prickles) than the other genotypes, which displayed low to no prickliness (Table 1).

#### 2.1.2 Reproductive traits

Regarding reproductive traits, substantial differences were observed among the genotypes (Table 2). Micro-Tom exhibited the earliest floral meristem development (D-FloralM), averaging 21.0 d post-transplantation, followed by Micro-Mel and ANG1 at 28.8 and 30.9 d, respectively, while Mini-Mel and MEL1 required 40.2 d. A similar trend was observed for flowering initiation (D-Flowering), although Mini-Mel flowered significantly earlier than MEL1. Specifically, Micro-Mel initiated flowering in 41.0 d, whereas Mini-Mel in 57.2 d (Table 2). Regarding the number of nodes to the first inflorescence (N-Nodes-Flower), Micro-Mel and Micro-Tom showed 7.9 and 8.0 nodes, respectively, followed by Mini-Mel and ANG1 with 9.4, while MEL1 had 10.2 nodes. ANG1, MEL1, and Mini-Mel produced few flowers per inflorescence (N-Flowers/Inf), with 2.1, 1.5, and 2.1 flowers, respectively. In contrast, Micro-Mel showed an average of 4.9 flowers, while Micro-Tom had an average of 9.3 flowers (Table 2).

**Table 2.** Mean values (± standard error) of the reproductive traits phenotyped for *S. anguivi* ANG1 and *S. melongena* MEL1 parental lines, Micro-Mel, Mini-Mel eggplant model varieties, and Micro-Tom tomato model variety. Ten plants of each genotype were evaluated. Traits reported in days were calculated from the date of seedling transplantation into pots (with a three-week interval between seed sowing and transplantation).

| Reproductive traits<sup>a</sup> | Genotypes |  |  |  |  |
| --- | --- | --- | --- | --- | --- |
|  | ANG1 | MEL1 | Micro-Mel | Mini-Mel | Micro-Tom |
| D-FloralM | 30.9 b ± 0.4 | 40.2 c ± 1.4 | 28.8 b ± 0.2 | 39.7 c ± 0.9 | 21.0 a ± 0.0 |
| D-Flowering | 43.2 b ± 0.5 | 60.2 d ± 1.8 | 41.0 b ± 0.6 | 57.2 c ± 0.8 | 27.9 a ± 0.4 |
| N-Nodes-Flower | 9.4 b ± 0.5 | 10.2 b ± 1.8 | 7.9 a ± 0.6 | 9.4 b ± 0.8 | 8.0 a ± 0.4 |
| N-Flowers/Inf | 2.1 a ± 0.1 | 1.5 a ± 0.2 | 4.9 b ± 0.5 | 2.1 a ± 0.3 | 9.3 c ± 0.4 |
| C-Colour | 3.0 a ± 0.5 | 7.0 b ± 1.8 | 7.0 b ± 0.6 | 9.0 c ± 0.8 | ND |
| D-FirstFruit | 52.4 b ± 1.4 | 72.5 d ± 1.6 | 48.6 b ± 0.5 | 63.4 c ± 1.2 | 36.0 a ± 0.3 |
| D-MaturityFruit | 87.0 b ± 3.0 | 123.2 e ± 2.7 | 103.7 c ± 1.9 | 114.7 d ± 1.7 | 71.4 a ± 0.6 |

<sup>a</sup>For each group, means (N=10) separated by different letters are significantly different at $p < 0.05$ according to the Student-Newman-Keuls parametric test.

Micro-Mel and MEL1 displayed light violet corollas (C-Colour), while Mini-Mel ones were bluish violet and those of ANG1 were white. In terms of fruit development, Micro-Tom was the earliest to set fruit at 36.0 d (D-FirstFruit), followed by Micro-Mel at 48.6 d, comparable to ANG1 at 52.4 d, then Mini-Mel at 63.4 d, and MEL1 the latest at 72.5 d (Table 2).

Maturing times (D-MaturityFruit) also followed a similar pattern, with Micro-Tom reaching the breaker stage at 71.4 d, followed by ANG1 at 87.0 d, Micro-Mel at 103.7 d, Mini-Mel at 114.7 d, and MEL1 at 123.2 d (Table 2). These data show that Micro-Mel enables 3.5 cycles per year under controlled growth chamber conditions, with an approximate duration of three and a half months from germination to fruit maturity. In contrast, Mini-Mel allows for an average of 3.0 cycles per year, requiring approximately four months per cycle. However, maturity time comparisons should be interpreted cautiously, as Micro-Tom and ANG1 may exhibit clearer colour transitions, whereas defining full maturity in eggplant is less precise due to subtler external ripening indicators.

#### 2.1.3 Fruit traits

The phenotypic evaluation of the fruits indicated that the largest fruits, i.e., exhibiting the greatest weight (Fruit-Weight), length (Fruit-Length), and width (Fruit-Width), were those of MEL1, followed by Micro-Mel (8.14 g weight, 27.51 mm length, 22.80 mm width) and Mini-Mel (6.92 g weight, 25.97 mm length, 21.43 mm width) (Table 3). Conversely, the ANG1 parental line consistently produced the smallest fruits. The fruit length/width ratio (Fruit-L/W-Ratio) showed the fruits of Micro-Tom and ANG1 were wider than longer, whereas those of MEL1, Micro-Mel, and Mini-Mel were longer than wider (Table 3). Regarding pedicel length, MEL1 displayed significantly the longest pedicel, followed by Micro-Mel (14.57 mm) and Mini-Mel (13.87 mm), ANG1, and Micro-Tom (Table 3).

**Table 3.** Mean values (± standard error) of the fruit traits phenotyped for for *S. anguivi* ANG1 and *S. melongena* MEL1 parental lines, Micro-Mel, Mini-Mel eggplant model varieties, and Micro-Tom tomato model variety. Ten plants of each genotype were evaluated.

| Fruit traits<sup>a</sup> | Genotypes |  |  |  |  |
| --- | --- | --- | --- | --- | --- |
|  | ANG1 | MEL1 | Micro-Mel | Mini-Mel | Micro-Tom |
| Fruit-Weight (g) | 0.59 a ± 0.03 | 29.87 d ± 5.44 | 8.14 c ± 0.92 | 6.92 c ± 1.04 | 3.96 b ± 0.13 |
| Fruit-Length (mm) | 8.06 a ± 0.15 | 42.32 d ± 3.45 | 27.51 c ± 1.60 | 25.97 c ± 1.79 | 16.36 b ± 0.18 |
| Fruit-Width (mm) | 9.71 a ± 0.21 | 35.07 d ± 2.95 | 22.80 c ± 0.84 | 21.43 bc ± 1.26 | 19.26 b ± 0.24 |
| Fruit-L/W-Ratio | 0.89 a ± 0.07 | 1.21 b ± 0.05 | 1.18 b ± 0.04 | 1.20 b ± 0.04 | 0.85 a ± 0.01 |
| Pedicel-Length (mm) | 9.95 b ± 0.16 | 39.60 d ± 0.87 | 14.57 c ± 0.43 | 13.87 c ± 0.45 | 4.00 a ± 0.05 |
| N-Seeds/Fruit | 28.2 a ± 2.1 | 107.3 b ± 36.0 | 46.3 a ± 10.5 | 19.7 a ± 9.9 | 22.9 a ± 1.2 |
| N-Seeds/Plant | 232.1 b ± 39.3 | 125.2 ab ± 36.9 | 211.1 b ± 38.1 | 43.8 a ± 23.0 | 351.9 c ± 23.7 |
| N-Fruits/Plant | 8.2 c ± 0.9 | 1.0 a ± 0.0 | 4.6 b ± 0.8 | 2.2 a ± 0.3 | 15.4 d ± 0.7 |
| Fruit-Curvature | 1.00 a ± 0.00 | 3.29 d ± 0.52 | 1.59 b ± 0.14 | 2.50 c ± 0.35 | 1.00 a ± 0.00 |
| Fruit-Shape | 5.00 a ± 0.00 | 5.00 a ± 0.00 | 5.63 b ± 0.15 | 4.90 a ± 0.10 | 5.00 a ± 0.00 |
| Apex-Shape | 4.24 a ± 0.17 | 7.00 c ± 0.00 | 5.71 b ± 0.26 | 4.30 a ± 0.36 | 4.77 a ± 0.07 |
| Calyx-Prickles | 0.00 a ± 0.00 | 1.00 c ± 0.00 | 0.02 a ± 0.02 | 0.50 b ± 0.11 | 0.00 a ± 0.00 |
| F-Colour-Phys-Mat | 5.95 b ± 0.03 | 2.00 a ± 0.00 | 2.07 a ± 0.04 | 2.00 a ± 0.00 | 5.93 b ± 0.02 |

<sup>a</sup>For each group, means (N=10) separated by different letters are significantly different at $p < 0.05$ according to the Student-Newman-Keuls parametric test.

MEL1 had the highest number of seeds per fruit (N-Seeds/Fruit), with 107.3 seeds/fruit on average, while there were no statistically significant differences for the rest of the genotypes, showing Micro-Mel a mean of 46.3 and Mini-Mel a mean of 19.7 seeds/fruit (Table 3). Micro-Tom produced the highest seed yield per plant (N-Seeds/Plant), with 351.9, followed by ANG1 (232.1), Micro-Mel (211.1), and MEL1 (125.2), with Mini-Mel yielding the fewest seeds per plant (43.8). In terms of fruits set per plant (N-Fruits/Plant), Micro-Tom had the highest value, with an average of 15.4 fruits/plant, followed by ANG1 (8.2) and Micro-Mel (4.6), whereas Mini-Mel and MEL1 produced a comparable number of fruits (2.2 and 1.0, respectively) (Table 3).

The analysis of fruit curvature (Fruit-Curvature) revealed that fruits of ANG1 and Micro-Tom were not curved, whereas those of Micro-Mel, Mini-Mel, and MEL1 exhibited a very slight curvature (Table 3). The fruit shape (Fruit-Shape) across all genotypes was substantially uniform, with the broadest section of the fruit located approximately halfway from the base to the tip, except for Micro-Mel, where it was slightly more proximal to the apex resulting in an ovoid shape (Table 3). The apex shape (Apex-Shape) varied from slightly protruded to depressed across the genotypes, with Micro-Tom, ANG1, and Mini-Mel showing slightly protruded apices, Micro-Mel having a more rounded apex, and MEL1 presenting a depressed apex. Regarding the presence of prickles on the calyx (Calyx-Prickles), Micro-Tom, ANG1, and Micro-Mel fruits were prickleless; Mini-Mel fruits displayed prickles in the calyx on half of the samples; and all MEL1 fruits had prickles in the calyx. Fruit colour at physiological maturity (F-Colour-Phys-Mat) was poppy red in Micro-Tom and ANG1 fruits, while it was deep yellow in MEL1, Micro-Mel, and Mini-Mel fruits (Table 3).

### 2.2 Regeneration assay

The *in vitro* regeneration evaluation revealed significant differences among the materials regarding shoot production, callus formation, and root development. The results highlight the distinct regeneration capacities observed in Micro-Mel and Mini-Mel compared to their parental lines. For the number of shoots per explant, MEL1, Micro-Mel, and Mini-Mel exhibited similar shoot production, with means of 1.2, 1.0, and 0.8 shoots per explant, respectively (Table 4). Conversely, ANG1 demonstrated a significantly higher shoot production compared to the other genotypes, averaging 4.6 shoots per explant, with its shoots showing a greater elongation (Figure 2). In terms of callus formation, Micro-Mel produced the highest number of calli, with an average of 1.4 cut edges with calli (Table 4). This was followed by MEL1 (1.3), Mini-Mel (1.2) and ANG1 (0.6). Surprisingly, the calli observed in Mini-Mel were purple due to the presence of anthocyanins (Figure 4G and 4H). Regarding root formation, an undesirable trait, MEL1 produced the highest number of roots, with an average of 0.1 roots per explant, which was significantly higher than all other genotypes (Table 4). In contrast, ANG1, Micro-Mel, and Mini-Mel exhibited minimal root formation, with an average of 0.0 roots per explant (Table 4).

**Table 4.** Mean values (± standard error) of the number of shoots, calli, and roots development during the regeneration test of *S. anguivi* ANG1 and *S. melongena* MEL1 parental lines, and Micro-Mel, Mini-Mel eggplant model varieties. A total of 600 explants, 150 per genotype, were evaluated.

| Regeneration traits<sup>a</sup> | Genotypes |  |  |  |
| --- | --- | --- | --- | --- |
|  | ANG1 | MEL1 | Micro-Mel | Mini-Mel |
| Number of shoots per explant | 4.6 b ± 0.2 | 1.2 a ± 0.1 | 1.0 a ± 0.1 | 0.8 a ± 0.1 |
| Number of cut edges with calli per explant | 0.6 a ± 0.1 | 1.3 b ± 0.0 | 1.4 c ± 0.0 | 1.2 b ± 0.1 |
| Number of roots per explant | 0.0 a ± 0.0 | 0.1 b ± 0.0 | 0.0 a ± 0.0 | 0.0 a ± 0.0 |

<sup>a</sup>For each group, means (N=150) separated by different letters are significantly different at $p < 0.05$ according to the Student-Newman-Keuls parametric test.

**Figure 2.**
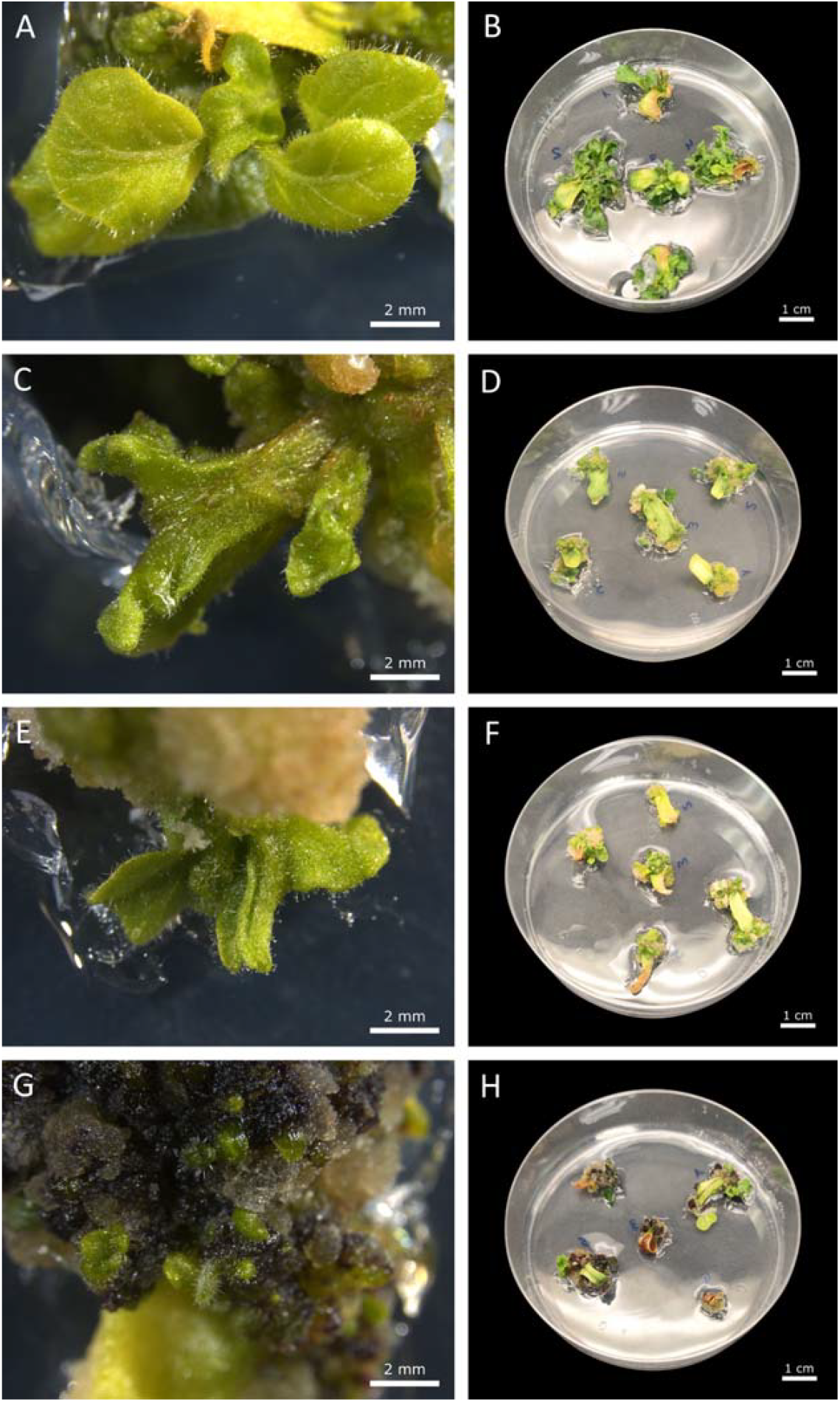
Shoots and callus formation in *S. anguivi* ANG1 (**A, B**), *S. melongena* MEL1 (**C, D**), Micro-Mel (**E, F**), and Mini-Mel (**G, H**) after one month of culture under a 16h light / 8h dark photoperiod. ANG1 showing well-elongated shoots with high production and minimal callus formation (A, B). MEL1 exhibiting moderate shoot production with compact calli (C, D). Micro-Mel displayed reduced shoot elongation and callus formation at the cut edges (E, F). Mini-Mel showed abundant purple-pigmented calli with limited shoot production (G, H). Images A, C, E, and G were taken with a Leica S Apo stereo microscope (Leica, Wetzlar, Germany).

**Figure 3.**
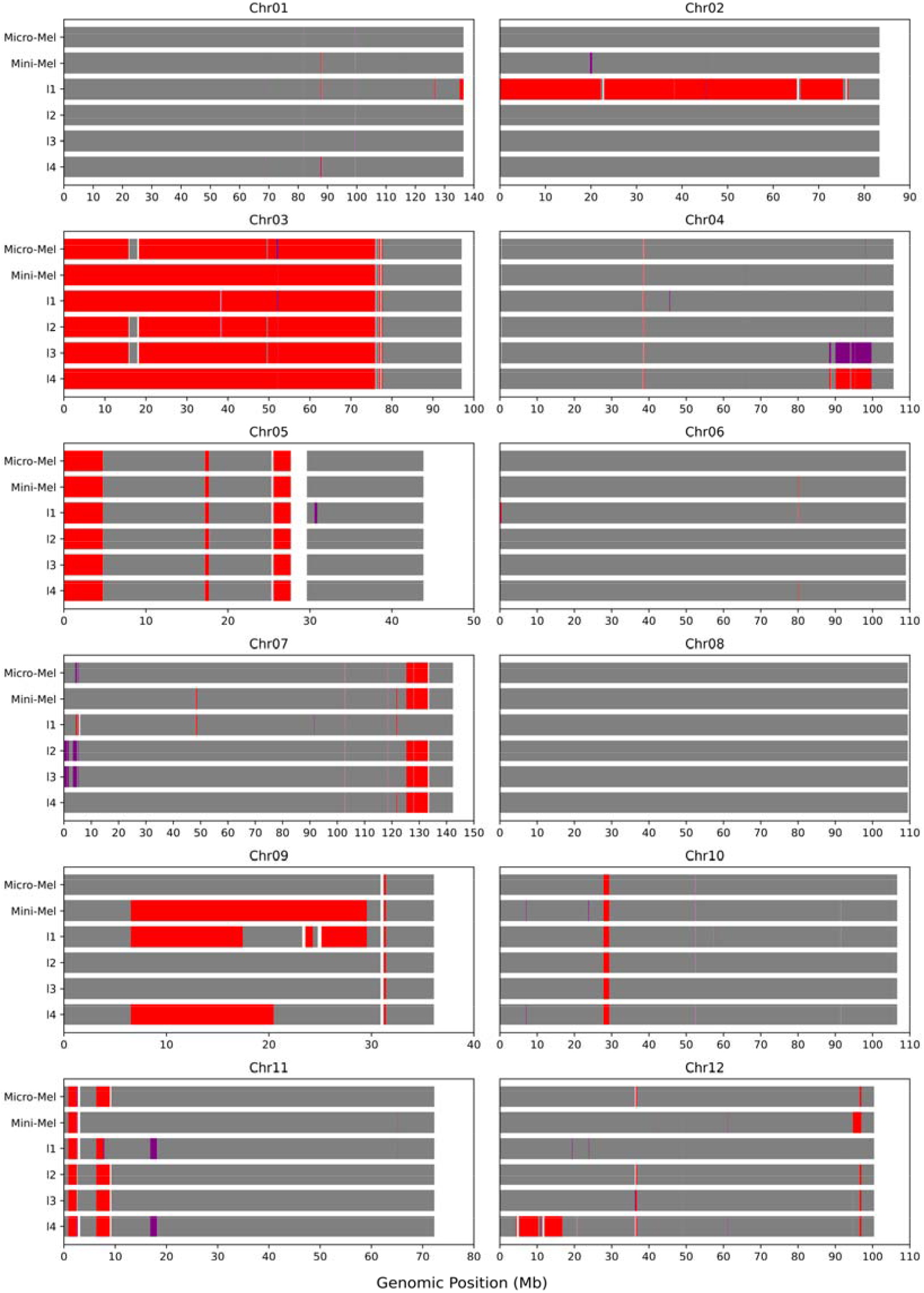
*Solanum anguivi* (ANG1) introgressed genomic regions in Micro-Mel, Mini-Mel, and four additional backcross dwarf materials (I1 to I4), all of them derived from the same original BC2 plant. The characterization of the introgressions was made with 179,653 high-quality SNPs between the parental lines identified by whole-genome sequencing. Homozygous introgressions are marked in red, and heterozygous introgressions are marked in purple.

**Figure 4.**
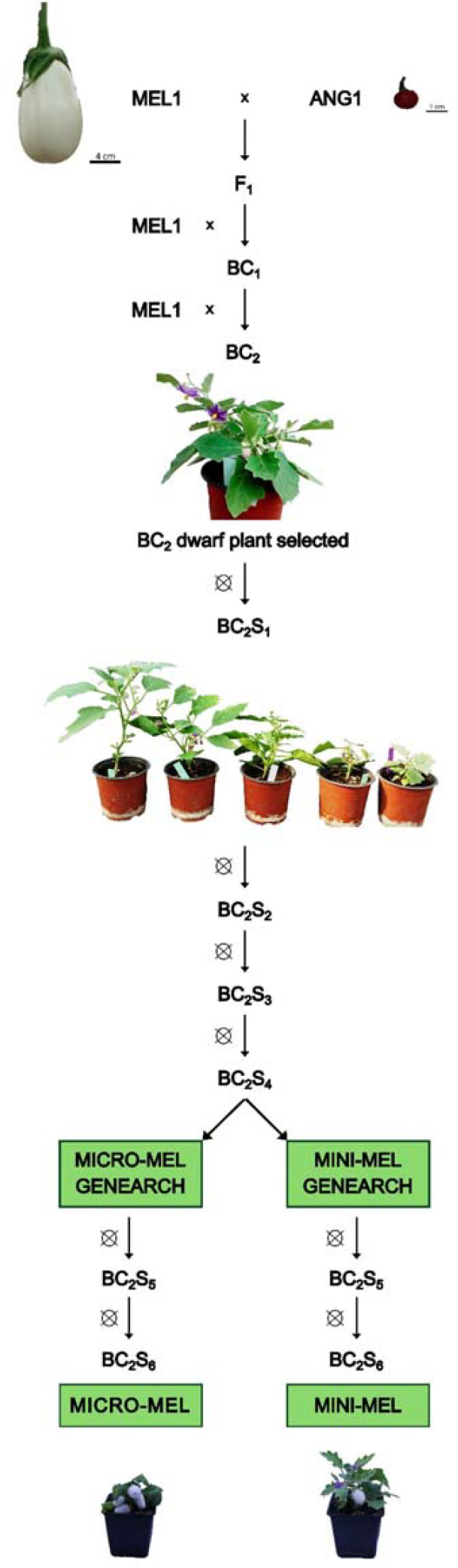
Breeding process for developing Micro-Mel and Mini-Mel eggplant model varieties. The process began with a cross between the two parental lines *S. melongena* (MEL1) and *S. anguivi* (ANG1), resulting in an F1 generation. The F1 was then backcrossed twice with MEL1 to generate a BC_2_ population. At this stage, the original dwarf phenotype was identified, selected, and self-pollinated to produce the BC_2_S1 generation. Subsequent self-pollination was conducted over four generations (BC_2_S1 to BC_2_S4), with phenotypic and genotypic selection using the SPET genotyping platform, resulting in BC_2_S4 lines. Within these lines, Micro-Mel and Mini-Mel were selected and self-pollinated up to the BC_2_S6 generation.

### 2.3 Genotyping, distribution and pattern of introgressions and candidate genes

Whole-genome sequencing of the two parental lines, along with Micro-Mel, Mini-Mel and the four intermediate BC_2_S_4_ materials, yielded an average of 188.73 M total clean reads per sample (Table S1). Overall, 97.02% of these bases had a quality score greater than 20 (Q20), indicating high sequencing accuracy. Mapping clean reads to the high-quality eggplant reference genome “67/3” v3.0 resulted in a 97.37% properly mapped read-pair rate, covering 69.85% of the reference genome with a mean depth of coverage of 23.37X (Table S1). After quality control filtering, a total of 179,653 SNPs were identified, with an average density of one SNP per 6,376 bp (Table S2). The average number of SNPs per chromosome was 14,971, ranging from 5,539 on chromosome 9 to 22,226 on chromosome 7 (Table S2). On average, 9,339 of these SNPs were heterozygous. Considering that the eggplant reference genome “67/3” v3.0 was covered by approximately 70.98% in Micro-Mel and 70.53% in Mini-Mel (Table S1), heterozygosity estimates of 0.00098% and 0.00123% were obtained for Micro-Mel and Mini-Mel, respectively (Table S3).

To identify the genomic localization of ANG1 introgressed genomic regions in the backcross dwarf lines, local ancestry analysis (LAI) implemented in Loter was performed [37], which reconstructs the mosaic of genomic segments inherited from the two parental lines. Genome-wide local ancestry patterns varied among the admixed individuals and exhibited heterogeneity along chromosomes (Figure 3, Table S4). On average, 10.95% of the ANG1 genome was introgressed into the backcross material, being 8.50% in Micro-Mel and 10.79% in Mini-Mel (Table S4). Micro-Mel contained 31 ANG1 recombinant blocks with an average length of 3.13 Mb, ranging from 17,211 bp on chromosome 4 to 31,175,003 bp on chromosome 3. On the other hand, Mini-Mel contained 41 ANG1 recombinant blocks with an average length of 3.01 Mb, ranging from 18,063 bp on chromosome 10 to 52,171,689 bp on chromosome 3 (Table S4). The resequencing data from the four additional dwarf backcross materials (I1 to I4), all derived from the same original BC2 plant, enabled the delimitation of potential genomic regions associated with the dwarf phenotype. As a result, 7.46% of the ANG1 genome was shared among the six introgressed materials, with its distribution detailed in Table 5. Annotation of these regions revealed 1,824 genes located within the shared ANG1 introgressions (Table 5). Chromosomes 3 and 5 contained the four longest consecutive regions of shared introgressions and, consequently, the highest number of genes inherited from ANG1 (Figure 3, Table 5). While no shared introgressions were found on chromosomes 2, 6, 8, and 12 (Figure 3, Table 5), the two haplotypes on chromosome 1, along with one on chromosome 4 and one on chromosome 7, were discarded as candidates for association with the dwarf phenotype, since no genes were found within those regions (Table 5).

**Table 5.** Shared *S. anguivi* ANG1 introgressed genomic regions among Micro-Mel, Mini-Mel, and four additional dwarf backcross materials (I1 to I4), all of them derived from the same original BC2 plant.

| Chromosome | Start (bp) | End (bp) | Length (bp) | Genes |
| --- | --- | --- | --- | --- |
| 1 | 81,841,398 | 81,879,851 | 38,453 | 0 |
|  | 99,428,057 | 99,472,127 | 44,070 | 0 |
| 3 | 8,748 | 15,773,894 | 15,765,146 | 573 |
|  | 18,283,553 | 75,963,141 | 57,679,588 | 762 |
|  | 76,872,818 | 77,149,486 | 276,668 | 19 |
|  | 77,468,548 | 77,723,534 | 254,986 | 17 |
| 4 | 38,617,662 | 38,782,274 | 164,612 | 2 |
|  | 65,801,459 | 65,818,670 | 23,894 | 0 |
|  | 98,137,428 | 98,216,814 | 79,386 | 4 |
| 5 | 52,195 | 4,758,459 | 4,706,264 | 379 |
|  | 17,209,221 | 17,708,071 | 498,850 | 12 |
|  | 25,551,183 | 27,661,523 | 2,110,340 | 28 |
| 7 | 102,849,361 | 102,913,969 | 64,608 | 2 |
|  | 118,495,672 | 118,551,791 | 56,119 | 0 |
| 9 | 31,194,173 | 31,413,894 | 219,721 | 3 |
| 10 | 27,790,391 | 29,369,925 | 1,579,534 | 4 |
|  | 105,776,693 | 105,794,756 | 18,063 | 2 |
| 11 | 827,344 | 2,512,177 | 1,684,833 | 17 |
| **TOTAL** |  |  | 85,265,135 | 1,824 |
| **TOTAL GENOME** |  |  | 1,142,799,992 |  |
| **PROPORTION (%)** |  |  | 7.46 |  |

A list of 30 genes linked to dwarfism in tomato, along with their respective orthologues in eggplant, is summarized in Table S5. These genes are distributed across all eggplant chromosomes, with the majority localized to chromosome 7 and only one situated on chromosome 9. Chromosomes 10, 11, and 8 also contain a notable number of orthologues (Table S5). The identified genes are involved in various processes related to plant growth and architecture, including gibberellin biosynthesis, cytokinin signaling, and cell wall formation (Table S5). Five genes associated with the dwarf phenotype in tomato, related to gibberellin biosynthesis and signaling (*GA2ox* and *DELLA*), have also been described in a recent eggplant dwarf mutant, along with two additional genes (Table S5). Within the shared introgressions from *S. anguivi* among the dwarf materials obtained in our work, three tomato orthologues were found: *SlERK1* (SMEL_003g176760.1.01, 36,249,765 - 36,255,417 pb) and *SlER* (SMEL_003g181050.1.01, 70,760,921 - 70,767,265 pb) on chromosome 3, and *SlDREB1* (SMEL_005g226710.1.01, 3,970,608 - 3,971,762 pb) on chromosome 5.

## 3. Discussion

Model plants are an essential tool in plant biology research enabling the study of genetic, physiological, and phenotypic processes in a more streamlined and manageable manner [15,38]. Here, we present two new model eggplant varieties, Micro-Mel and Mini-Mel, which share key traits such as compact size, short internodes, fast growth, multiple inflorescences, small white fruits and short life cycle. However, they differ in key aspects that broaden their applications. Micro-Mel has determinate growth, an earlier flowering and fruit set time than Mini-Mel, which makes it particularly suitable for studies requiring rapid generation turnover.

In comparison, Mini-Mel, with its indeterminate growth, greater number of internodes, and thicker stem, provides an alternative model for morphological and physiological studies or those requiring a more extended vegetative phase. These distinctive characteristics enable efficient use of space in large-scale studies, allowing for experiments across multiple generations within a shorter time frame. Micro-Mel allows 3.5 cycles per year under controlled growth chamber conditions and Mini-Mel an average of 3.0 cycles per year. This is especially beneficial for advancing genetic studies and plant breeding programs by significantly reducing the typical long growth cycles in eggplant [39,40]. Additionally, the ability to cultivate them in a climatic chamber reduces the dependence on external weather conditions and minimizes environmental variability [41,42].

Although other reduced-sized eggplant lines have been reported [3,43], none exhibit the unique combination of dwarfism and short life cycle characteristics of Micro-Mel and Mini-Mel. They offer unique opportunities for advancing eggplant research and provide practical advantages for urban agriculture and space-efficient cultivation systems, particularly in small-scale agricultural settings [8]. Furthermore, they adhere to the Different, Uniform, and Stable (DUS) criteria [44,45], ensuring high uniformity across individuals (estimated heterozygosity of 0.00093% for Micro-Mel and 0.00117% for Mini-Mel) and reliable preservation of their defining characteristics over successive generations. Moreover, unlike Micro-Tom, which has multiple versions due to the lack of genomic tools available at the time of its development [15], Micro-Mel and Mini-Mel have been precisely characterized both phenotypically and genomically. Their genetic composition has been defined accurately through whole-genome sequencing, ensuring a clear and standardized reference for future research. Meeting these criteria reinforces their potential as model plants and underscores their value for breeding programs and experimental reproducibility in plant research.

The Micro-Mel and Mini-Mel varieties hold significant promise as model plants for eggplant research, similar to the role that Micro-Tom has played in tomato studies. Micro-Tom has contributed extensively to tomato research by enabling high-density cultivation and short life cycles, facilitating studies in genetics, genomics, and breeding [8,14,15]. Likewise, Micro-Mel and Mini-Mel offer similar advantages for eggplant research, allowing faster high-throughput phenotyping and genetic studies in a space-efficient manner. Pathogen resistance studies have also benefited from Micro-Tom’s short life cycle and compact size [46], suggesting that Micro-Mel and Mini-Mel could facilitate controlled pathogen assays in eggplant, particularly for quarantine diseases or those requiring prolonged incubation periods. Additionally, Micro-Tom has been useful in studying plant responses to abiotic stress [12,13], an application where Micro-Mel and Mini-Mel could be valuable due to their adaptability to controlled environments, enabling standardized drought, salinity, and temperature stress experiments. In gene editing research, Micro-Tom has provided a robust system for CRISPR-based modifications [20,21,47], which could be replicated in Micro-Mel and Mini-Mel to accelerate gene function studies in eggplant. Furthermore, Micro-Tom has supported reverse genetics approaches such as TILLING and mutant library development [48,49], indicating that Micro-Mel and Mini-Mel could be used to establish similar resources for the eggplant research community. Just as near-isogenic lines (NILs) have been developed in Micro-Tom to study the effects of specific alleles on fruit traits [50], similar approaches could be applied to Micro-Mel and Mini-Mel. For example, their white fruits provide an ideal background for generating NILs that differ only in fruit colour, facilitating the precise evaluation of fruit pigmentation-related genes.

Beyond their value as research tools, Micro-Mel and Mini-Mel also exhibit strong potential for ornamental horticulture. Similar to dwarf tomatoes, compact peppers, and miniature cabbages [29,31–33,35,36], their compact size and growth habit make them well-suited for decorative use. Their suitability for cultivation in small gardens, balconies, or indoor spaces enhances their versatility beyond research applications. Additionally, their short internodes, lobed leaves, and vivid violet flowers, combined with the production of small white/yellow fruits, contribute to their ornamental appeal, positioning them as promising candidates for ornamental markets.

Efficient *in vitro* regeneration is a critical attribute for any model plant system, as it enables gene editing and functional studies [20,21]. The regeneration results highlight the potential of Micro-Mel and Mini-Mel as model plants, as both lines regenerate like the cultivated parental MEL1, producing approximately one shoot per explant on average. Both dwarf lines exhibited reliable regeneration, albeit with lower efficiency than reported in other studies. For instance, Yesmin et al. (2021) reported a mean regeneration efficiency ranging from 1.87 to 4.70 shoots per explant, depending on the medium and genotype [25], while García-Fortea et al. (2020) achieved an average of 9 shoots per explant using the same explant type and medium as those employed in this study [23]. In comparison, some studies on Micro-Tom reported a mean of 15.8 shoots per cotyledon explant [51], and 2-3 shoots per explant under a different medium formulation [52]. Although the regeneration efficiency observed in this study is lower than these reports, the results remain highly encouraging and might be improved by optimizing media compositions based on protocols that have demonstrated higher success rates [25,53,54]. In this way, Micro-Mel and Mini-Mel proved consistent and reliable regeneration capabilities, comparable to the cultivated parental MEL1, underscoring their potential as robust model plants. Their regeneration capacity could significantly streamline and expedite genetic transformation and gene editing experiments, rendering these lines invaluable for both fundamental and applied research in eggplant.

In addition to the morphological characterization of Micro-Mel and Mini-Mel, this study identified 179,653 polymorphic SNPs between MEL1 and ANG1 to obtain a comprehensive representation of their mosaic genomes. Both compact eggplant model varieties shared ANG1 haplotypes on chromosomes 1, 3, 4, 5, 7, 9, 10, 11, and 12, but notable differences in the introgressed regions were observed, which may underlie the previously described phenotypic differences. A major difference was observed on chromosome 9, where Mini-Mel carried a large ANG1 introgression spanning 6,5196,644 to 29,557,107 bp, which was absent in Micro-Mel. To identify genomic regions associated with the dwarf phenotype, resequencing data from four additional backcross dwarf materials enabled the identification of 18 shared introgressions. Our results suggest that the mechanism underlying dwarfism in these materials could involve several possibilities. First, the presence of orthologues of tomato genes linked to dwarfism, *SlDREB1* [55], *SlER* and *SlSERK1* [8], within the shared introgressions points to modifications in the gibberellin pathway [55–57] or in the interaction between the *SlER* gene and its receptor *SlSERK1*, both involved in the regulation of stem length [8,58,59]. Second, the presence of additional shared introgressions among the backcross materials could indicate that the genetic basis of dwarfism in eggplant may differ significantly from that in tomato, with the introgressions from *S. anguivi* potentially introducing novel mechanisms not previously identified in tomato [60–62]. Finally, the dwarf phenotype observed in Micro-Mel and Mini-Mel could also be due to epistatic interactions between genes introgressed from *S. anguivi* and the genetic background of *S. melongena* [63–66]. These results highlight the complex genetic network contributing to the dwarf phenotype in eggplant and emphasize the need for further research to elucidate the specific mechanisms involved. Ongoing backcrosses to the *S. melongena* parental line are aimed at refining our understanding of these genetic pathways.

## 4. Conclusion

The development of Micro-Mel and Mini-Mel represents a significant breakthrough in the creation of model plants for eggplant research. These compact and fast-growing short life cycle lines combine the essential traits of model systems (space efficiency, rapid growth cycles, and reliable regeneration) with distinct morphological characteristics that make them uniquely suited for diverse research and horticultural applications. Their ability to adhere to DUS criteria ensures consistency and reproducibility, reinforcing their utility in genetic, physiological, and phenotypic studies. Moreover, Micro-Mel exceptional compactness and accelerated life cycle position it as a cornerstone for advancing genetic studies and breeding programs, while Mini-Mel morphology offers versatility for studies requiring a more conventional plant architecture. The ornamental potential of both lines further expands their appeal beyond research, offering practical and aesthetic value for urban and small-scale horticulture. By providing an innovative platform for eggplant studies, Micro-Mel and Mini-Mel are poised to become highly valuable tools in plant biology and crop improvement, closing the gap between eggplant and other important Solanaceae species in terms of research capabilities and applications.

## 5. Materials and methods

### 5.1. Plant material

Micro-Mel and Mini-Mel are derived from the cross between the white-fruited common eggplant *S. melongena* (MEL1) and the eggplant wild relative *S. anguivi* (ANG1) [67]. Both accessions are conserved in the germplasm bank of COMAV at Universitat Politècnica de València, València, Spain. The interspecific MEL1 x ANG1 hybrid was backcrossed (BC) towards MEL1 and 20 plants of the BC_1_ generation were individually backcrossed again to MEL1 to obtain the BC_2_ (Figure 4). Among the BC_2_ plants, one potted plant displayed a compact size, semi-determinate growth, and profuse fruiting of small white fruits. The observations were confirmed by transplanting the original plant outdoors, displaying consistent features.

The selected BC_2_ dwarf plant was self-fertilized (Figure 4), and progeny selection was based on the desired ideotype, characterized by reduced leaf and flower size, compact growth with short internodes and prolificacy (high set of seeded fruits). The preselected BC_2_S_1_ plants were genotyped to identify those with fewer introgressions from *S. anguivi*. For this, young leaf tissue was sampled, and genomic DNA was extracted following the SILEX protocol [68]. DNA quality and integrity were evaluated via agarose electrophoresis and NanoDrop ND-1000 spectrophotometer (NanoDrop Technologies, Wilmington, Delaware, USA) while DNA concentration was estimated using a Qubit® 2.0 fluorometer (Thermo Fisher Scientific, Waltham, MA, USA). The samples were then sent to Identity Governance and Administration (IGA) Technology Services (IGATech, Udine, Italy) for library preparation and sequencing on a NextSeq500 sequencer (150 paired-end) for a high-throughput genotyping with the 5k probes SPET eggplant platform [69]. Sequence analysis was performed using the Trait Analysis by Association, Evolution, and Linkage (TASSEL) software (ver. 5.0) [70].

The selected BC_2_S_1_ plants were self-fertilized and this process was repeated until the BC_2_S_4_ generation (Figure 4). At this stage, the genearchs of Micro-Mel and Mini-Mel were selected and further stabilized through two additional rounds of self-pollinations (BC_2_S6) using a pedigree approach, where the final Micro-Mel and Mini-Mel lines were established. The entire development process of Micro-Mel and Mini-Mel, involved over 300 plants and spanned nine generations (Figure 4). To ensure the ideotype stability, the plants were grown in pots of various sizes and under different environmental conditions, including climatic chambers, glasshouses, and open field, confirming their high phenotypic stability (Figure 5). In addition to Micro-Mel and Mini-Mel, their two parental lines (MEL1 and ANG1), the subsequent intermediate generations, and the tomato Micro-Tom variety was also phenotyped for comparison.

**Figure 5.**
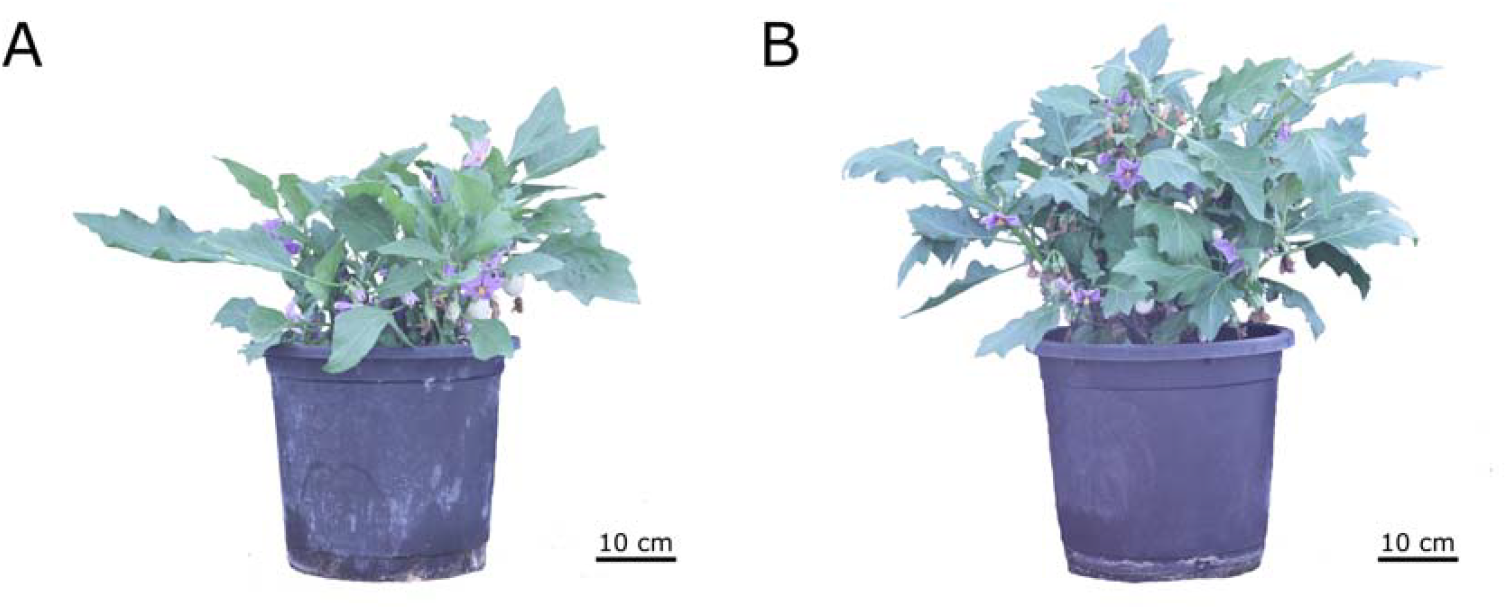
Micro-Mel (**A**) and Mini-Mel (**B**) eggplant model plants cultivated in a large pot (18 L), maintaining their dwarf phenotype.

### 5.2 Phenotypic characterization of Micro-Mel and Mini-Mel

Seed germination was conducted *in vitro* following a modified germination protocol developed by [71] to overcome the seed dormancy typically observed in the wild species. Seeds of ANG1, MEL1, Micro-Mel, and Mini-Mel were sterilized through immersion in 70% ethanol for 30 s, followed by immersion in 20% commercial bleach with 3.8% NaOCl for 15 min. Subsequently, the seeds were washed three times in sterilized water for 3 min. All seeds were then incubated for one day in a sterilized solution of gibberellic acid 3 (GA3) at 500 mg/L.

Seeds were sown *in vitro* in Microbox containers (O119/140+OD119/140 #10 (G), SACO2, Nevele, Belgium) containing a base medium consisting of 4.4 g/L of Murashige and Skoog (MS) medium [72], 20 g/L of sucrose, 1 mg/L of calcium pantothenate, 0.1 mg/L of biotin, and 9 g/L of MicroAgar. Seedlings were cultured *in vitro* under a photoperiod of 16 h light/8 h dark at a constant temperature of 25°C. After two weeks, seedlings were transplanted into seedbeds with Humin substrate N3 (Klasmann-Deilmann, Germany). One week later, 10 plants of each genotype were transferred to 11×11×12 cm pots (1.5 L), with five plants randomly distributed per tray within the growth chamber. The trays were moved every week to reduce the environmental effect within the climatic chamber. Uniform watering was provided as needed.

Thirty morphological traits were measured, including traits listed in the eggplant descriptors of UPOV (2020) and IBPGR (1990). These traits were classified into three categories: vegetative, reproductive, and fruit traits (Table 6). Vegetative and reproductive traits were measured on individual plants within each genotype, while fruit traits were assessed on several individual fruits to calculate a representative mean for each plant.

**Table 6.** List of evaluated traits including abbreviations and assessment methods. Traits reported in days were calculated from the date of seedling transplantation into pots (after a three-week interval between seed sowing and transplantation).

| Traits | Abbreviation | Assessment |
| --- | --- | --- |
| **Vegetative** |  |  |
| Plant height at 30 days | P-Height-30 | From the base to the apical growing point (cm) |
| Plant height at 60 days | P-Height-60 | From the base to the apical growing point (cm) |
| Plant height at 90 days | P-Height-90 | From the base to the apical growing point (cm) |
| Plant height at 120 days | P-Height-120 | From the base to the apical growing point (cm) |
| Stem width at 30 days | S-Width-30 | At the lower stem in contact with the soil (mm) |
| Stem width at 60 days | S-Width-60 | At the lower stem in contact with the soil (mm) |
| Stem width at 90 days | S-Width-90 | At the lower stem in contact with the soil (mm) |
| Stem width at 120 days | S-Width-120 | At the lower stem in contact with the soil (mm) |
| Number of internodes at 60 days | N-Internodes | From the base to the apical growing point |
| Length of internodes at 60 days | L-Internodes | cm |
| Growth habit | Growth-Habit | 3: upright, 5: intermediate, 7: prostrate |
| Growth type | Growth-Type | 0: indeterminate, 1: determinate |
| Anthocyanin presence in the upper leaves at 90 days | Anthoc-UL | 0: absence, 1: presence |
| Anthocyanin presence in the stem | Anthoc-Stem | 0: absence, 1: presence |
| Leaf blade lobing | Leaf-Lobing | 1: very weak or none, 3: weak, 5: intermediate, 7: strong, 9: very strong |
| Foliar prickles presence | Foliar-Prickles | 0: absence, 1: presence |
| **Reproductive** |  |  |
| Number of days to the appearance of the floral meristem | D-FloralM | First developing flower bud visible on the main stem |
| Number of days to flowering | D-Flowering | First open flower |
| Number of nodes to the first inflorescence | N-Nodes-Flower | - |
| Number of flowers per inflorescence | N-Flowers/Inf | - |
| Corolla colour | C-Colour | 1: greenish white, 3: white, 5: pale violet, 7: light violet, 9: bluish violet |
| Number of days to the appearance of the first fruit | D-FirstFruit | Increase in size in the first ovary and pedicel |
| Number of days to the physiological maturity of the first fruit | D-MaturityFruit | First fruit showing an intense and uniform colour |
| **Fruit** |  |  |
| Fruit weight | Fruit-Weight | Including the pedicel (g) |
| Fruit length | Fruit-Length | From the apex of the fruit to the receptacle (mm) |
| Fruit width | Fruit-Width | Maximum fruit width (mm) |
| Fruit length/width ratio | Fruit-L/W-Ratio | Fruit length/fruit width |
| Pedicel length | Pedicel-Length | From the point of attachment to the stem to the point of attachment to the calyx (mm) |
| Number of seeds per fruit | N-Seeds/Fruit | - |
| Number of seeds per plant | N-Seeds/Plant | - |
| Number of fruits per plant | N-Fruits/Plant | - |
| Fruit curvature | Fruit-Curvature | 1: none/fruit straight, 3: slightly curved, 5: curved, 7: snake-shaped, 8: sickle-shaped, 9: u shaped |
| Fruit shape | Fruit-Shape | Position of the widest part of the fruit: 3: about 1/4 way from base to tip, 5: about 1 /2 way from base to tip, 7: about 3/4 way from base to tip |
| Apex shape | Apex-Shape | 3: protruded, 5: rounded, 7: depressed |
| Prickles presence in calyx | Calyx-Prickles | 0: absence, 1: presence |
| Fruit colour at physiological maturity | F-Colour-Phys-Mat | 1: green, 2: deep yellow, 3: yellow-orange, 4: deep orange, 5: fire red, 6: poppy red, 7: scarlet red, 8: light brown, 9: black |

### 5.3 In vitro regeneration assay

For the *in vitro* regeneration assay, a total of three replicates were conducted. Each replicate included 50 explants for Micro-Mel, Mini-Mel, and their parental lines MEL1 and ANG1, with five explants per plate. This setup resulted in a total of 120 plates, encompassing 600 explants, 150 per genotype. The seed sterilization was performed as described before, except for the germination medium used. The seeds were sown in the E0 medium formulation described by García-Fortea et al. (2020), which consists of 2.2 g/L MS basal medium [72], 15 g/L sucrose, and 7 g/L gelrite [23]. The seeds were kept in a climatic chamber at 25°C under a 16 h light/8 h dark photoperiod. Twelve days after sowing, the proximal and distal portions of the cotyledons were excised, and the explants were then placed abaxial side down onto Petri dishes containing the eggplant regeneration medium E6 [23]. The regeneration medium consisted of 2.2 g/L MS medium, 15 g/L sucrose, 7 g/L gelrite, and 2 mg/L zeatin riboside (ZR). After one month of cultivation, the number of shoots per explant, cut edges with calli, and roots per explant were measured. Images were captured using a Leica S Apo stereo microscope (Leica, Wetzlar, Germany).

### 5.4 Statistical analysis

Means and standard errors were calculated for phenotyping and organogenic response data for the genotypes evaluated. Differences among the genotypes were evaluated with a one-way ANOVA. The significance of differences was assessed with a Student-Newman-Keuls multiple range test at a *p* < 0.05 significance level. The statistical data analysis was performed using Statgraphics Centurion 19 software (Statgraphics Technologies, Inc., The Plains, VA, USA).

### 5.4 Whole-genome sequencing

Seeds from the two parental lines MEL1 and ANG1, Micro-Mel, Mini-Mel, and four intermediate BC_2_S_4_ materials (l1, l2, l3 and l4), were germinated *in vitro* and genomic DNA was extracted as previously described. High-quality DNA samples (260/280 and 260/230 ratios > 1.8) were sent to the Beijing Genomics Institute (BGI Genomics, Hong Kong, China) for the construction of 150 bp paired-end libraries and subsequent whole-genome sequencing (WGS) on the DNBseq platform. Raw sequence data were deposited in the NCBI Sequence Read Archive (SRA) under the Bioproject identifier PRJNA1230165.

Raw reads underwent quality control using fastq-mcf v.1.04.676 [73] with a quality threshold of 30 (−q 30) and a minimum remaining sequence length of 50 bp (−l 50). High-quality reads were then aligned against the high-quality eggplant reference genome “67/3” v3.0 [74] using the BWA-MEM algorithm v. 0.7.17–r1188 with default parameters [75]. PCR duplicates were removed using Picard MarkDuplicates tool v. 1.119 (https://broadinstitute.github.io/picard/). Variant calling was performed using Freebayes v. 1.3.6 [76] with default configuration settings, except for the minimum mapping and base quality thresholds set to 20. Candidate SNPs underwent three filtering rounds. The first round, conducted with VCFtools v. 0.1.16 (https://vcftools.github.io/man_latest.html), involved selecting sites with an allele frequency greater than 0.1 (−-maf 0.1) and no missing data (−-max-missing 1), and genotypes supported by at least 10 reads (−-minDP 10). The second round, performed with BCFtools v. 1.13 (https://samtools.github.io/bcftools/bcftools.html), retained biallelic SNPs with distinct genotypes between the parental lines. In the final step, PLINK v1.9.0-b.7.7 [77] was used to prune the SNP data based on linkage disequilibrium (LD) with the parameter ‘--indep-pairwise 50 5 0.7’. Heterozygosity was estimated by calculating the proportion of heterozygous genotypes in the final set of high-quality SNPs, obtained using VCFtools v. 0.1.16 (−-het), relative to the total sequenced genome.

### 5.6 Identification of introgressions and search for gene candidates

To detect ANG1 introgressions in the genetic backgrounds of Micro-Mel, Mini-Mel, and the four intermediate BC_2_S_4_ materials, the command-line implementation of Loter was used [37], a non-parametric software for local ancestry inference. This software assumes that haplotypes in admixed individuals originate from hybridization and recombination between reference parental genomes [37]. Before ancestry inference, phasing was performed on the high-confidence SNP set obtained after three filtering rounds using Beagle 5.2 [78]. The phased reference dataset included genotypes from the donor and recurrent parental lines, while the admixed dataset consisted of phased genotypes from the six backcross dwarf materials. Identified introgressions were subsequently filtered, removing those supported by fewer than five variants. Finally, results were visualized using Python 3.9.21 matplotlib v. 3.9.2 package [79]. Once the shared introgressions among the six backcross dwarf materials were identified, the search for gene candidates was conducted through a comprehensive literature review. The primaly focus was to identify genes that have been described as potential candidates for dwarfism in eggplant, as well as in tomato, the closest species with extensive genomic information and well-established synteny with eggplant. The tomato orthologues were searched in the eggplant reference genome “67/3” v3.0 using the BLAST tool from the Sol Genomics Network (https://solgenomics.sgn.cornell.edu/tools/blast/?db_id=303).

## Acknowledgments

This work was funded from grant PID2021-128148OB-I00, funded by MCIN/AEI/10.13039/501100011033/ and “ESF Investing in your future” and from grant PDC2022-133513-I00 funded by MICIU/AEI/10.13039/501100011033 and by the European Union NextGeneration EU/PRTR. Funding was also received from grant CIPROM/2021/020 funded by Conselleria d’Educació, Universitats i Ocupació of the Generalitat Valenciana. MM-L has received a predoctoral grant (FPU21/02288) funded by the Spanish Ministerio de Ciencia, Innovación y Universidades. VB-F has received a predoctoral grant (CIACIF/2023/238) founded by Conselleria d’Educació, Cultura, Universitats i Ocupació (Generalitat Valenciana). PG has received a postdoctoral grant (RYC2021-031999-I) funded by MICIU/AEI/10.13039/501100011033 and by the European Union NextGeneration EU/PRTR.

## Author contributions

### M.M.-L

Conceptualization, Data curation, Formal analysis, Investigation, Methodology, Validation, Visualization, Writing - Original Draft. *V.B.-F*.: Data curation, Formal analysis, Investigation, Methodology, Software, Writing - Original Draft. *E.G.-F*.: Investigation, Resources, Writing - Review and Editing. M.P.: Funding acquisition, Project administration, Resources, Writing - Review and Editing. *S.V*.: Funding acquisition, Project administration, Resources, Writing - Review and Editing. J.P.: Conceptualization, Funding acquisition, Methodology, Project administration, Resources, Supervision, Writing - Review and Editing.

### P.G

Conceptualization, Methodology, Project administration, Resources, Supervision, Validation, Writing - Review and Editing. All authors have read and agreed to the published version of the manuscript.

## Data availability statement

The documentation for the registration of Micro-Mel and Mini-Mel as protected varieties is currently being prepared to apply for plant breeders’ rights protection through the community plant variety office (CPVO). Seeds of both materials are freely available for research and educational studies. Commercial uses will require a licensing agreement. Requests for seeds and commercial use can be directed to the corresponding author.

Raw sequence data in this study can be found in the NCBI Sequence Read Archive (SRA) under the Bioproject identifier PRJNA1230165.

## Supplementary information

Supplementary information is available at *Horticulture Research* online.

