## Supplementary Figures for "Micro-Mel and Mini-Mel short life cycle dwarf lines with *Solanum anguivi* introgressions as the first model varieties for eggplant research and breeding"

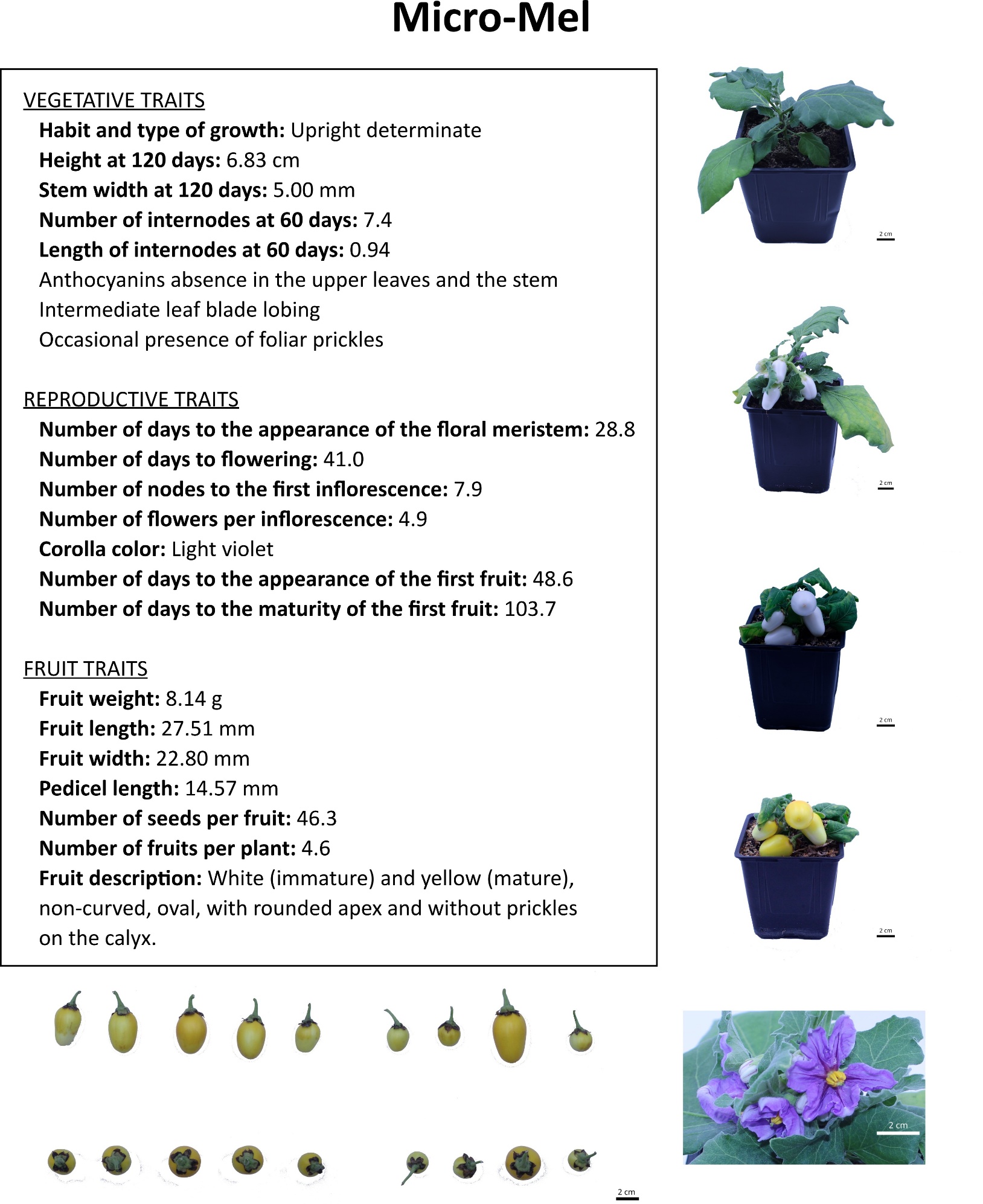

**Figure S1.** Variety fact sheet for the Micro-Mel dwarf eggplant variety.

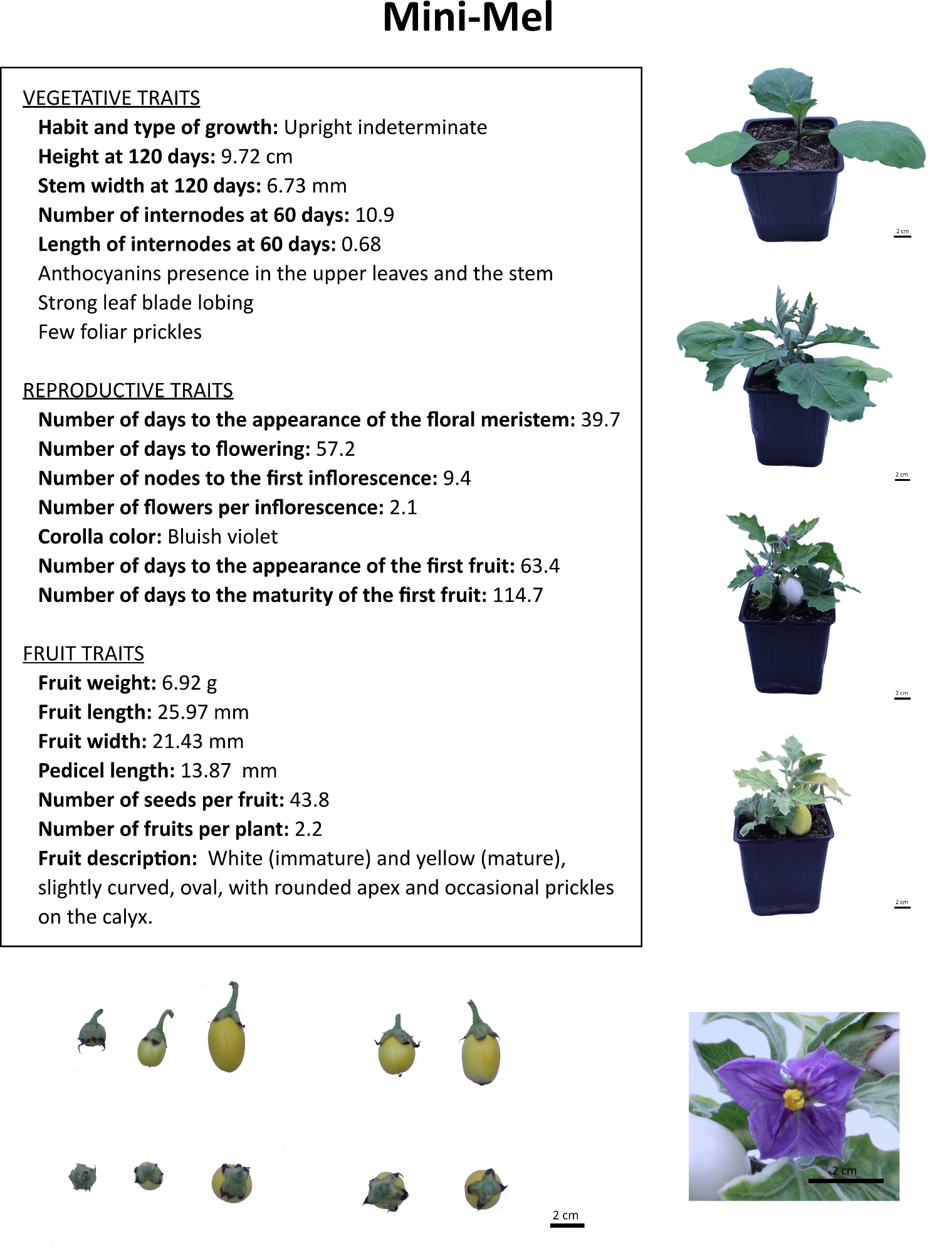

**Figure S2.** Variety fact sheet for the Mini-Mel dwarf eggplant variety.
